# Agency improves working memory and accelerates visual and attentional processing

**DOI:** 10.1101/2020.10.22.350397

**Authors:** Rocio Loyola-Navarro, Cristóbal Moënne-Loccoz, Rodrigo C. Vergara, Alexandre Hyafil, Francisco Aboitiz, Pedro E. Maldonado

**Author notes:** Corresponding author: Pedro E. Maldonado, Ph.D., Departamento de Neurociencia, Facultad de Medicina, Universidad de Chile, Independencia 1027, Santiago, Chile.

## Abstract

Agency, understood as the ability of an organism to control stimuli onset, modulates perceptual and attentional functions. Since stimulus encoding is an essential component of working memory (WM), we conjectured that the perceptual process’s agency would positively modulate WM. To corroborate this proposition, we tested twenty-five healthy subjects in a modified-Sternberg WM task under three stimuli presentation conditions: an unpredictable presentation of encoding stimulus, self-initiated presentation of the stimulus, and self-initiation presentation with random-delay stimulus onset. Concurrently, we recorded the subjects’ electroencephalographic signals during WM encoding. We found that the self-initiation condition was associated with better WM accuracy, and earlier latencies of N100 and P200 evoked potential components representing visual and attentional processes, respectively. Our work demonstrates that agency enhances WM performance and accelerates early visual and attentional processes deployed during WM encoding. We also found that self-initiation presentation correlates with an increased attentional state compared to the other two conditions, suggesting a role for temporal stimuli predictability. Our study remarks on the relevance of agency in sensory and attentional processing for WM.

## INTRODUCTION

While classic Working Memory (WM) paradigms require subjects to passively attend to task-relevant stimuli, that are presented without control of the participants, in everyday situations, stimuli relevant to WM tasks may appear to our sensory system as a consequence of the agent’s movements, such as what occurs during eye movements or when we click on the mouse or the mobile screen to browse a web page. When the agent initiates the presentation of the stimuli, behavior and its underlying neural mechanisms are modulated in several cognitive processes such as perception (Blakemore et al. 1999) and attention (Nittono et al. 2003a; Nittono 2004; Nittono 2006; Waszak & Herwig 2007; Kumar et al. 2015). This idea seems to be in line with the embodied cognition theory (O’Regan & Noë 2001; Varela et al. 2016), which states that subjects’ bodies, particularly their motor systems, influence cognition. Nevertheless, it remains unknown whether agency, understood as the control of stimuli onset by the agent (i.e., self-initiation), modulates WM encoding processing. Additionally, it has not been established if this influence is achieved by modulating sensory, attentional processes or later updating mechanisms of stimuli in memory. Revealing possible motor modulations of WM through self-initiation of stimuli may improve our understanding of the WM mechanism in real-life situations, hallmarking participants’ role as agents of their own cognitive process.

What are the mechanisms deployed in agency that can possibly modulate WM encoding? Evidence in perceptual and attentional tasks have shown that the agents’ self-initiation of stimuli modulates both sensory and attentional neural processing (Nittono et al. 2003b; Walsh & Haggard 2008; Stenner et al. 2014; Concha-Miranda et al. 2019). This finding is relevant in WM’s context since WM has been proved to rely on both perceptual (Teng & Kravitz 2019; Zhang et al. 2019) and attentional processing (Postle 2006). Furthermore, attention is proposed as a critical component of WM, being responsible for the maintenance of items and the WM span limits (Cowan 2005). One proposed mechanism by which agency may influence perception is the modulation that movements exert on early sensory cortices (Schroeder et al. 2010; Stenner et al. 2015; Concha-Miranda et al. 2019). It has been reported in animal models that motor acts such as eye movements (specifically saccades) modulate primary sensory cortex firing rate and local field potentials (Ito et al. 2011; McFarland et al. 2015). In humans, a similar modulation has been described in the P100 Event Related Potential (ERP) component, which has been proposed as an index of the early visual response in the visual cortex (Luck et al. 1990; Clark et al. 1995; Di Russo et al. 2002; Sergent et al. 2005; Natale et al. 2006). Devia et al. (2017) reported that the amplitude of the P100 component is larger when the stimulus was fixated after a saccade compared to flashed stimuli (i.e., non-saccade-mediated fixation) in a free-viewing paradigm. They also described earlier latencies of the P100 component, suggesting that the visual cortex activates faster to visual stimuli when they are a consequence of a motor act, such as saccades, compared to visual stimuli that are passively sensed (Devia, Montefusco-Siegmund, Egaña, Maldonado 2017). The motor cortex activity, such as the supplementary motor area (SMA), has been proved to modulate the visual cortex in mice through anatomically direct connections (Leinweber et al. 2017). Although this has not been confirmed in humans, a hand movement such as a button press could also modulate the cortical visual processing through SMA-visual cortex connections. If this is true, self-initiation of the stimulus during WM encoding should yield a modulation in the early visual encoding indexes such as P100 and N100 ERP components. A similar mechanism could be operating in the modulation observed of self-initiation on attentional neural substrates. The evidence in attentional tasks have shown that agency increases the amplitude of both P300a and P300b components compared to passive externally paced stimuli (Nittono et al. 2003a; Nittono 2004; Nittono 2006). This evidence reveals that agency enhances attentional processing through both attentional capture (indexed by P300a) and updating mechanisms in memory (indexed by P300b). These findings suggest that motor commands could also modulate WM at a later stage of encoding. SMA is also connected to the cortical regions related to the attentional mechanisms involved in WM, namely posterior parietal cortex (PPC) (Reep et al. 1994; Wilber et al. 2015; Bozkurt et al. 2017) and dorsolateral prefrontal cortex (DLPFC) (Hasan et al. 2013; Bozkurt et al. 2017). Suppose SMA is modulating the activity of the PPC and DLPFC. In that case, a self-initiated encoding should be associated with a modulation of attention indexes during WM encoding, such as P300 (Polich 2007) and P200 (Dunn et al. 1998; Missonnier et al. 2004) ERPs components. There are no current reports about self-initiation modulation on the P200 component; however, earlier P200 latency correlates with better performance in WM tasks (Missonnier et al. 2007). As for motor actions modulating WM, self-initiation has been explored (McMillan et al. 2007; Holgado et al. 2019), but it has never been directly compared to passive conditions. There is, instead, behavioral evidence stating that task-unrelated voluntary motor acts worsen WM outcome (Gündüz et al. 2017), suggesting that motor sequence execution shares cognitive resources with WM. Nevertheless, whether this interference results from a sensory or/and an attentional neural modulation remains unknown.

Self-initiated tasks usually allow for temporal prediction of the stimuli appearance, since the movement that triggers the stimuli is finely coupled in time (i.e., time-locked) to the sensory consequence. Time-prediction, even if it does not occur in the context of self-initiation, refers to a neural mechanism in which the “processing and detection of events are facilitated by minimizing the uncertainty about when they are going to occur’’ (Arnal & Giraud 2012). If the agency’s modulatory effects rely on time-prediction only, then the jittering of the motor act and its sensory consequence should abolish those effects. Fine temporal coupling between motor acts and their sensory consequences have been depicted to be crucial to visual cortex development (Attinger et al. 2017) and important in perception (Blakemore et al. 1999) and attention (Morillon et al. 2014). However, studies indicate that movements that are made before and after stimuli presentation (hence, the stimuli are not an immediate consequence of the motor act) can also influence perception (Tomassini et al. 2015) and even long-term memory performance (Yebra et al. 2019).

This evidence is suggestive of a possible motor-related modulation that is independent of temporal prediction. Yebra et al. (2019) indicated that the noradrenergic system’s engagement mediates this motor-related improvement in long-term memory performance during encoding. Since the noradrenergic system massively modulates brain activity, it is possible to speculate an influence at both early sensory and attentional encoding processing levels.

Based on the literature reviewed above, we hypothesized that the agents’ self-initiation of stimuli modulates WM encoding. This effect relies on the temporal modulation of visual, attentional and memory updating processing. To test this hypothesis, 25 healthy adults performed a modified Sternberg WM task (Sternberg 1966), with three different ways of deploying the stimuli, while EEG activity was recorded. The classic control WM paradigm (Passive (P) condition), consisting of the automatic presentation of the stimuli, was compared to two self-initiated conditions: an active coupled condition (AC) and an active decoupled condition (AD). If active self-initiated WM encoding improves WM through a temporal modulation of visual, attentional or memory updating processing, we should find: a) better performance in AC compared to P, b) an effect in encoding the ERPs markers in AC compared to P, and c) a relation between ERP modulation and performance. Moreover, to test if agency influence is based on temporal prediction, a decoupled encoding condition (AD) was designed. AD consisted of the presentation of the stimuli after a random delay (400-600 ms) after the button press, which reduces the temporal predictability of the stimuli onset without affecting the sense of agency (Wen 2019). In case the modulatory effects of agency rely on time-prediction only, we should find: a) no significant differences in performance and ERP components between AD and P conditions, since both conditions lack of precise time-prediction of stimuli onset, and b) statistically significant difference in performance and ERP components between AC and AD. Our results show an effect of task conditions on both performance and ERP components, with AC yielding better performance and earlier latencies of ERP components compared to both AD and P conditions. Moreover, AD condition also presents better performance and earlier latencies of ERP components compared to P condition, suggesting that agency does not rely on the temporal coupling between action and its sensory consequence only.

## MATERIALS AND METHODS

### Participants

Twenty-six healthy adults (13 females; mean age 23.1 y.o.; range 18-31 y.o.) volunteered and were tested. All the participants were under- or postgraduate students of the Universidad de Chile with non-current or history of neurological, psychiatric, systemic disease, and normal or corrected-to-normal vision. This information was corroborated by anamnesis and the application of the MoCA test (Nasreddine et al. 2005). In the MoCA test, participants had to score ≥ 26 points to be included. As for anamnesis, the exclusion criteria were: a) Cranioencephalic trauma; b) Usage of illegal drugs during the last three months; c) Uncompensated systemic disease (metabolic disease or epilepsy); d) Usage of one or more of these drugs: benzodiazepines, anticonvulsants, metilfenidate, modafinil; and e) Diagnosis of depression or adult attentional deficit disorder. Informed consent was previously read and signed by the participants. One of the 26 participants was excluded because of luminance differences during testing. Consent for the current study was approved by the Ethical Committee in Human’s Research of the Medicine Faculty of the Universidad de Chile.

### Task

Participants engaged in a modified Sternberg working memory task (Sternberg 1966), with three encoding conditions while recording EEG activity. The three conditions differed only in how the stimuli were triggered (whether controlled by the agent or externally) while sharing the same time course from stimuli presentation until the participants’ answers. Conditions corresponded to an Active Coupled condition (AC), an Active Decoupled condition (AD) and a Passive condition (P). Details of the task scheme are shown in Figure 1.

**Figure 1.**
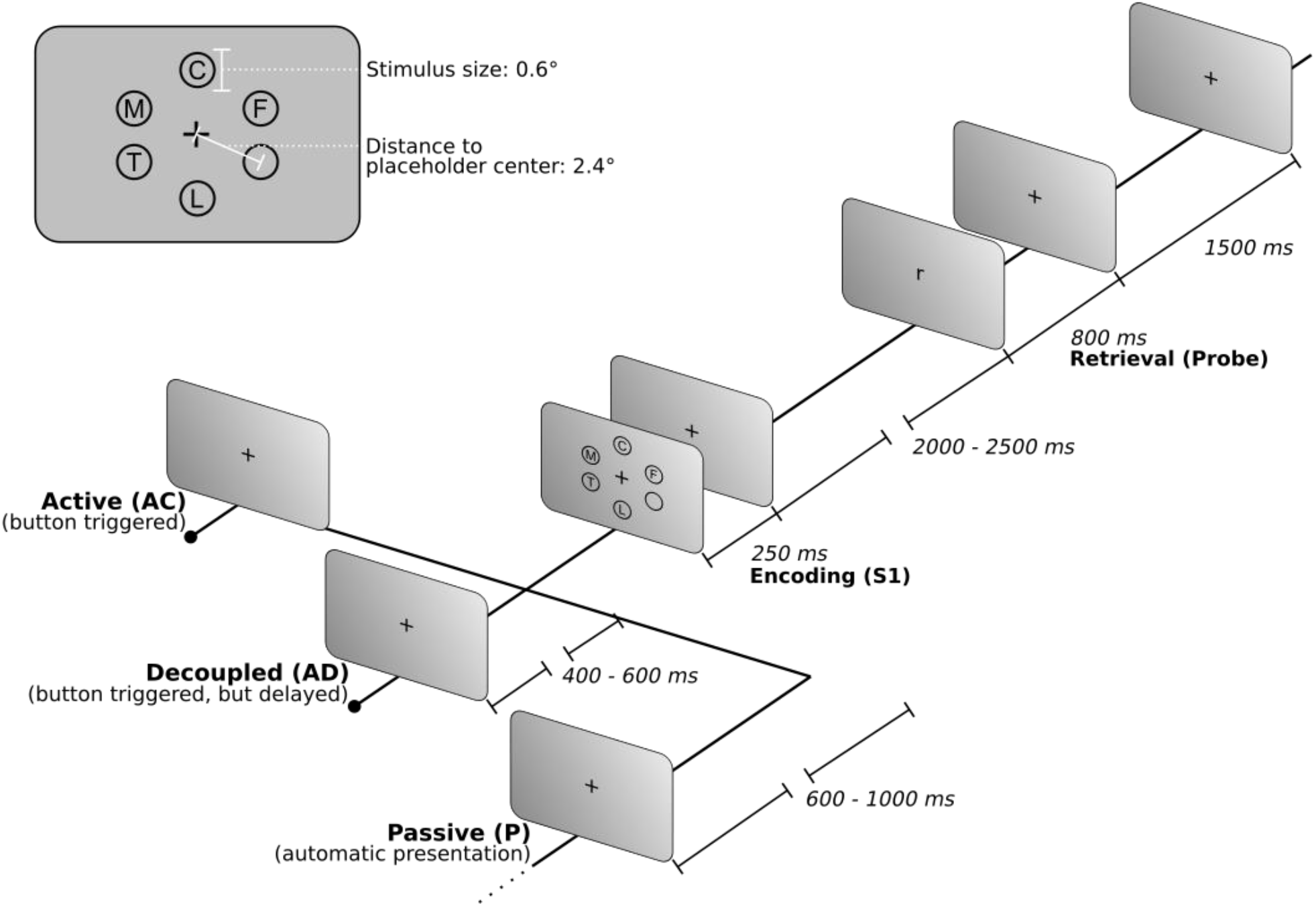
Schematic presentation of the WM task showing the three experimental conditions. Each condition varies only on the pre-encoding stage. Active coupled condition (AC) starts with a button press, and the stimuli array (S1) is displayed immediately. Decoupled condition (AD) also starts with a button press, but the S1 is delayed by a random time between 400 and 600 ms. Finally, Passive condition (P) follows an automatic presentation of S1 at a random time between 600 and 1000 ms following fixation cross onset. The inset shows an example of the S1 array and its visual disposition. (° visual degrees; ms: milliseconds).

In all the three conditions, each trial began with an eye-tracking-based drift correction to ensure that participants’ eyes positions remain similar in every trial. After drift-correction, a first fixation cross marked the beginning of each trial. The stimuli array (S1) lasted 250 milliseconds (ms). After S1 disappeared, only the central fixation cross remained for a random time between 2000 and 2500 ms (maintaining). Subsequently, the probe stimulus appeared for 800 ms, followed by the final fixation cross, which lasted 1500 ms (retrieval). Subjects had all the retrieval time to respond (800 ms + 1500 ms = 2300 ms). If the probe was present in the S1 array, participants had to press the joystick’s right back button with their right index finger. Conversely, if the probe was absent from set S1, participants had to press the left-back button with their left index finger.

As stated before, to determine if the agents’ self-initiation of WM modulates WM, the onset of the S1 set was manipulated (i.e., the pre-encoding stage), generating three task conditions: AC, AD, and P. In the AC condition, participants were required to press a frontal button of the joystick (with either of their thumbs). At the same time, the first fixation cross was present. Immediately after the button press, the S1 array appeared on the screen for 250 ms. The AD condition was very similar: participants were also required to press a frontal button. At the same time, the first fixation cross was present, but in this condition, S1 appeared with a random delay (400-600 ms) after the button press. This time-lapse does not disturb the sense of agency of individuals (Wen 2019). The P condition corresponds to the classic automatic presentation of the stimuli, which appear after a random fixation time of 600-1000 ms.

Each condition was presented in separate blocks of 100 trials. Each participant executed all the three blocks, accomplishing a total of 300 trials. The presentation order of the blocks was pseudo-randomized per participant. Instructions were given separately for each condition. Participants were not told about the specific difference between AC and AD conditions to not bias their answers. For participants 12 to 25, at the end of each block, they were asked to determine the task’s difficulty. To do so, they rated their global sensation of how secure they made their answers in that particular block (hereafter “confidence”). A scale of confidence ranging from 1 to 7 was used, with 1 corresponding to “not sure of my answers,” 4 being neutral, and 7 corresponding to “very sure of my answers.” The experimental paradigm was designed in Experiment Builder (SR Research Ltd., Mississauga, Ontario, Canada) and executed in Eyelink 1000 (SR Research Ltd., Mississauga, Ontario, Canada).

### Materials

The stimuli were presented in a flat *Viewsonic* 27 inches screen (23.54×13.24 inches; 1920×1080 pixels of resolution). Subjects were seated at a 72 cm distance from the screen. Therefore, 40.35 pixels equals 1 visual degree (°). Images were 45.1° wide and 26.3° high. Subjects were required to maintain their chin on a chinrest during the whole experiment to reduce head movements and maintain a stable distance between the eyes and the screen. Fatigue blinks and eye positions were monitored with an Eyelink 1000 (SR Research Ltd., Mississauga, Ontario, Canada) eye-tracking system.

The background was set to gray (127/127/127 RGB) during the whole session to avoid luminance changes. All the stimuli (S1 and probe stimulus) were black consonants. S1 consisted of 5 capital consonants (except X, Y) presented simultaneously in a circular array surrounding a fixation cross (Fig. 1). The fixation cross appeared at the center of the screen (at pixel coordinates 960,540). Stimuli positions were demarcated by six circles as placeholders (based on Proksch & Bavelier 2002; Green & Bavelier 2003). Therefore, one circle was left empty in every trial. Each of the six circle positions had the same probability of being empty. As for the stimuli’s size, consonants were set to a height of 0.6°, while the placeholders’ circles had 1° diameter (based on Proksch & Bavelier 2002; Green & Bavelier 2003). Placeholders centroids were set at a 2.4° distance from the central fixation cross. Since the parafoveal region has a size of 5.2° (Kolb 1995), this disposition ensured that the stimuli inside the placeholders fit into the parafoveal region. Three hundred S1 sets were created, one for each trial. Each S1 set consisted of five unrepeated consonants chosen randomly by “rand” function in Matlab©. Consonants “B”, “V”, and “W” never appeared in the same set S1; the same occurred with “M” and “N”. This was done to avoid acoustical or visual associations. As for the probe stimulus, it consisted of a unique consonant (excluding “x” or “y”) presented in the center of the screen (coordinates 962,540). The presence or absence of the probe stimuli on the S1 set had the same probability (50%). Responses were made by pressing a button with either left (probe is not present on S1) or right (probe is present in S1) index finger. A joystick, model Microsoft SideWinder, was used to record the response.

### Behavioral data analysis

To assess the effects of agents’ self-initiation on WM, the encoding condition’s effect on accuracy and reaction times was analyzed by performing two One-way-repeated-Analysis of Variance (ANOVA) and pertinent post hoc tests when needed. Accuracy was defined as the proportion of correct answers (True Positives and True Negatives) for each participant. Reaction time (RT) corresponded to the time (in ms) elapsed since the appearance of S2 until the answer made with the button press. The effects of the encoding condition on the two dependent variables were analyzed using two One-way-repeated ANOVA. Post hoc tests for each variable with significant effects were computed using a paired-t-test corrected by the Holm-Sidak method’s multiple comparisons. These parametric tests were chosen according to results of normality distribution (assessed by Shapiro-Wilk test) and of equality of variances (assessed by Levene test) of both accuracy and RT (See Table 1).

**Table 1.**
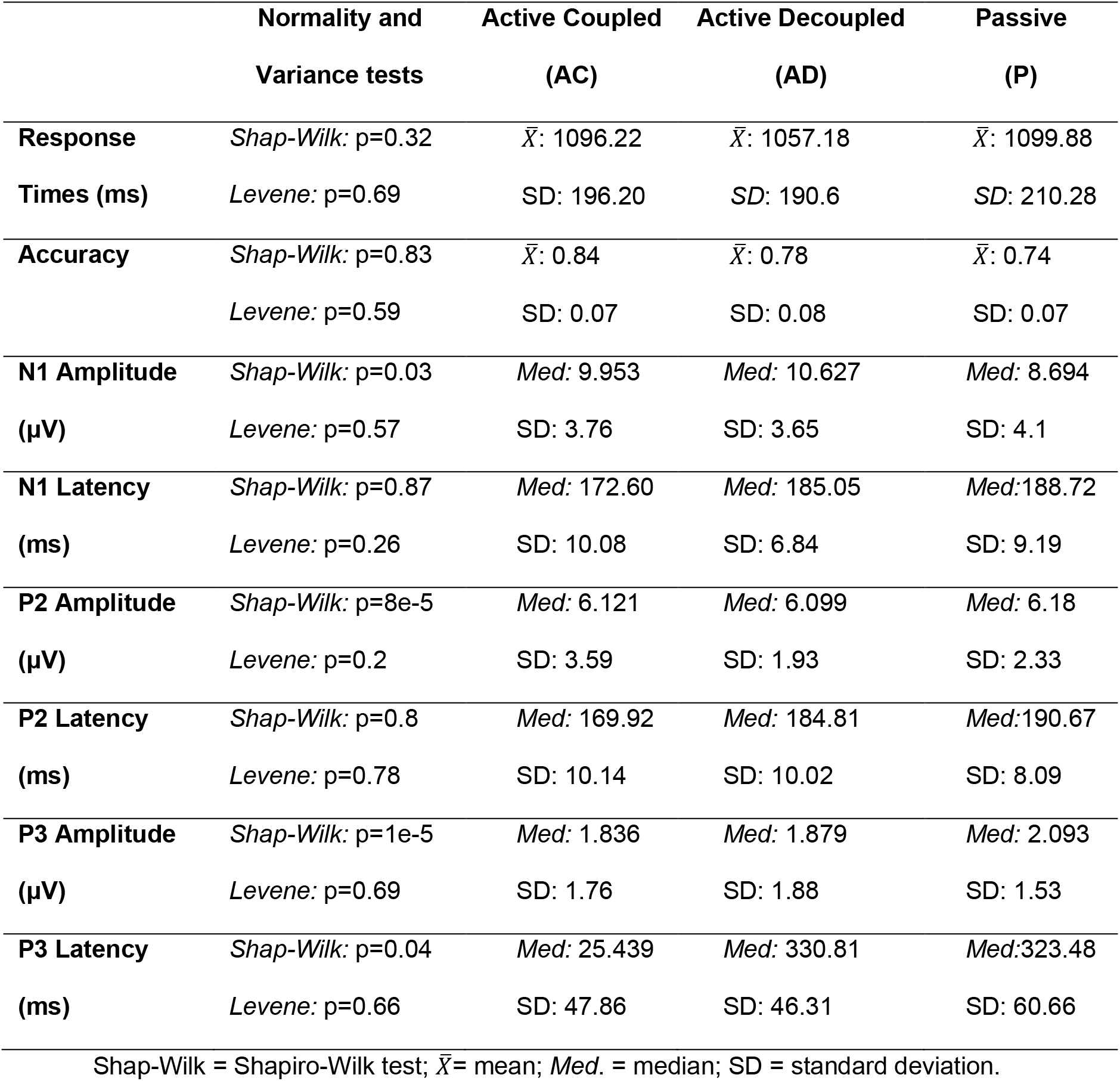
Values of the dependent and independent variables per encoding condition

The effects of other variables that could influence WM behavior were tested next. Thus, a binomial General Linear Mixed Model (GLMM) was performed (Moscatelli et al. 2012). Trial accuracy was set as the dependent variable and participant as a random effect. GLMM included the following variables as fixed effects: encoding condition, task learning, block position, and prestimuli time. Task learning was defined as the change in the probability of making a correct response due to the increasing trial number (from 1 to 100). The block position was a categorical variable, and its value could be first, second, or third block. Finally, pre-stimuli time was analyzed in two different ways (and therefore, in two separate models): a) a total pre-stimuli time was defined as the time elapsed from the first fixation cross onset until the onset of the S1 array, b) uncontrolled pre-stimuli time, defined as the time previous to stimuli onset that the subjects could not control. In the passive condition, both the total and uncontrolled pre-stimuli times were the same: the first fixation cross duration. In the AD condition, uncontrolled pre-stimuli time was the delay time between the button press and the S1 set appearance; total pre-stimuli time was the delay time plus the time the participant took to press the frontal button while the first fixation cross was present. Finally, in the AC condition, the uncontrolled pre-stimuli time is 0, while the total prestimuli time was the time the participant took to press the frontal button. All the four independent variables were standardized using z-score transformation. The Akaike information criterion (AIC) was applied to select the best model.

### Electroencephalography and Signal pre-processing

Electroencephalographic (EEG) activity was recorded at a 2048 Hz sample rate using a BioSemi Inc. amplifier of 32 active scalp electrodes and eight external electrodes to record eye movements. Common Mode Sense (CMS) and Driven Right Leg (DRL) electrodes were used as ground electrodes. Head caps were utilized to hold electrodes according to the 10/10 system (Jurcak et al. 2007). Eight external electrodes were set: one for each mastoid and six to record the eye movements (EOG): three around the right orbit and three around the left orbit.

Continuous EEG signal was re-referenced offline to the average activity of the 32 electrodes. Signal was then filtered using a FIR symmetric passband filter between 0.5 and 40 Hz with a linear phase. Its design is firwin with a Hamming window (acausal, zero-phase delay, and one-pass forward). The size of the filter was 6.6 seconds. The transition bandwidth of the filter was 0.5 Hz in the lower frequency limit and 10 Hz in the upper-frequency limit. The passband ripple of the filter was 0.0194 dB and the stopband attenuation was 53 dB.

Noisy channels were eliminated by visual inspection and then interpolated using spherical splines. After that, an Independent Component Analysis (ICA) was performed to determine and eliminate components related to blinks and eye movements. Segments containing muscle artifacts and other artifacts unrelated to blinks were eliminated automatically through a 500 μV peak-to-peak rejection threshold.

### Event-Related Potential (ERP) Components calculation

Continuous Signal was divided into epochs centered on the stimuli’s appearance. The epoch was set to 500 ms before and 1000 ms after the stimuli appearance. Noisy epochs were rejected using a 250 μV peak-to-peak threshold. We explored ERP components related to those processes to analyze the influence of agency in early visual, attentional, and memory updating processing. Components of interest were *N100-like, P200-like, and P300-like* (hereafter, N1, P2, P3). We used electrode Oz to calculate P1 and N1 components, Fz to calculate the P2 component, and Pz to obtain the P3 component. All ERPs’ components were computed by averaging correct cleaned trials. Amplitudes and latencies of each ERP component were assessed. Amplitude was calculated using peak-to-peak values. We used this approach since a component near 0 ms appeared in AC condition only, which could influence the peak amplitude values in the AC condition. The peak-to-peak method consisted of calculating the component’s peak amplitude in a certain electrode and time-window, then calculating a reference value and subtracting both the values. To determine each component’s peak value, the latency of that component was first calculated from the grand-average (average of the ERP across all 25 subjects). The peak amplitude and latency in each subject were then calculated in a 100 ms window around the grandaverage latency. The reference value for the N1 component corresponded to P1, while the reference values for P2 and P3 were defined as the negative peak appearing immediately after either P2 or P3. This data was calculated using only 24 of the 25 subjects since one of the participants was discarded because he had no reliable ERP responses. EEG data were preprocessed and averaged using MNE-Python (Gramfort 2013).

### Event-Related Potential (ERP) statistical analysis

To test if there is an effect of the encoding condition over amplitude and latency, Kruskal-Wallis, or ANOVA tests (one per ERP component) were performed, designed as CONDITIONSXLATENCIES and CONDITIONSXAMPLITUDES. The Rank-sum Wilcoxon or t-test corrected by Holm-Sidak was performed as a Post Hoc test. These tests were selected based on the distribution of the data (amplitude and latency) using the Shapiro-Wilk and Levene tests, respectively. To assess which electrophysiological activity distinguishes better among experimental conditions, a conditional Classification Tree (CART model) was used. CART models allow us to classify or estimate phenomena with discrete changes (Hothorn, Hornik, Van De Wiel, et al. 2006; Hothorn, Hornik, & Zeileis 2006; Strobl et al. 2009). For this particular CART model, the Monte Carlo Method was selected, with 1000 resampling to estimate tree splits’ significance using an alpha of 0.05. Additionally, to avoid overfitting the model, the amount of observation per leaf was limited to 20% of the total observations. The partitioning variables used to characterize the experimental conditions were accuracy, RT, and the amplitudes and latencies of P1N1, P2, and P3 components.

To test if the relationship between the better electrophysiological variable and accuracy is dependent on the task condition, a Linear Mixed Model was performed. The model included accuracy as a dependent variable, the best electrophysiological variable (yielded by the CART model) as a fixed effect, and the encoding conditions as a random effect variable.

All computations were performed using RStudio (Rstudio Team 2016) for statistical computing. Libraries used were EZ (v4.4-0) (Lawrence 2016), car (v3.0-2) (Fox & Weisberg 2014), party (v1.3-5) (Hothorn et al. 2015), and lme4 (v1.1-21) (Bates et al. 2015).

## RESULTS

The current work hypothesized that the self-initiated WM stimulus presentation improves WM performance through the temporal modulation of visual, attentional, and memory updating processing. To evaluate how the agent’s self-initiation of stimuli would affect WM encoding, 25 participants performed a Sternberg working memory task, manipulating the stimuli’s onset. In the Active Coupled (AC) condition, the stimuli were presented immediately after the participants pressed a button. In the Active Decoupled condition (AD), stimuli were presented with a delay of 400 to 600 ms after the button press, which allowed to test whether possible effects of self-initiation are tied to temporal precision of stimulus onset. Finally, these two self-initiated conditions were contrasted with the passive presentation of the stimuli (P), where stimuli were automatically presented between 600-1000 ms after the first fixation onset. Each encoding condition consisted of one block of 100 trials so that each subject executed three separate blocks (one per encoding condition) of 100 trials each (See <u>Materials and Methods</u>).

Two approaches tested the hypothesis proposed: i) Is behavior modulated by these three experimental conditions? and ii) Does self-initiation modulate early visual, attentional, and memory updating processing during encoding?

### 1. Agents’ self-initiation of the stimuli enhance WM performance

To explore whether the agency modulates WM mechanisms, we tested the effect of Encoding Conditions (AC, AD, and P) on performance (RT and accuracy). Table 1 shows averages, standard deviations, and results of Shapiro-Wilk and Levene tests for reaction times (RT) and accuracy, specified per encoding condition. The ANOVA results reveal a main effect of Encoding condition on accuracy (F(2,48) = 25.67, p = 2.6e-08; η^2^G= 0.2). No Encoding Condition effects were found on RT (F(2,48) = 1.06, p = 0.35) (Fig. 2). These results reveal that agency modulates accuracy without affecting RT.

**Figure 2.**
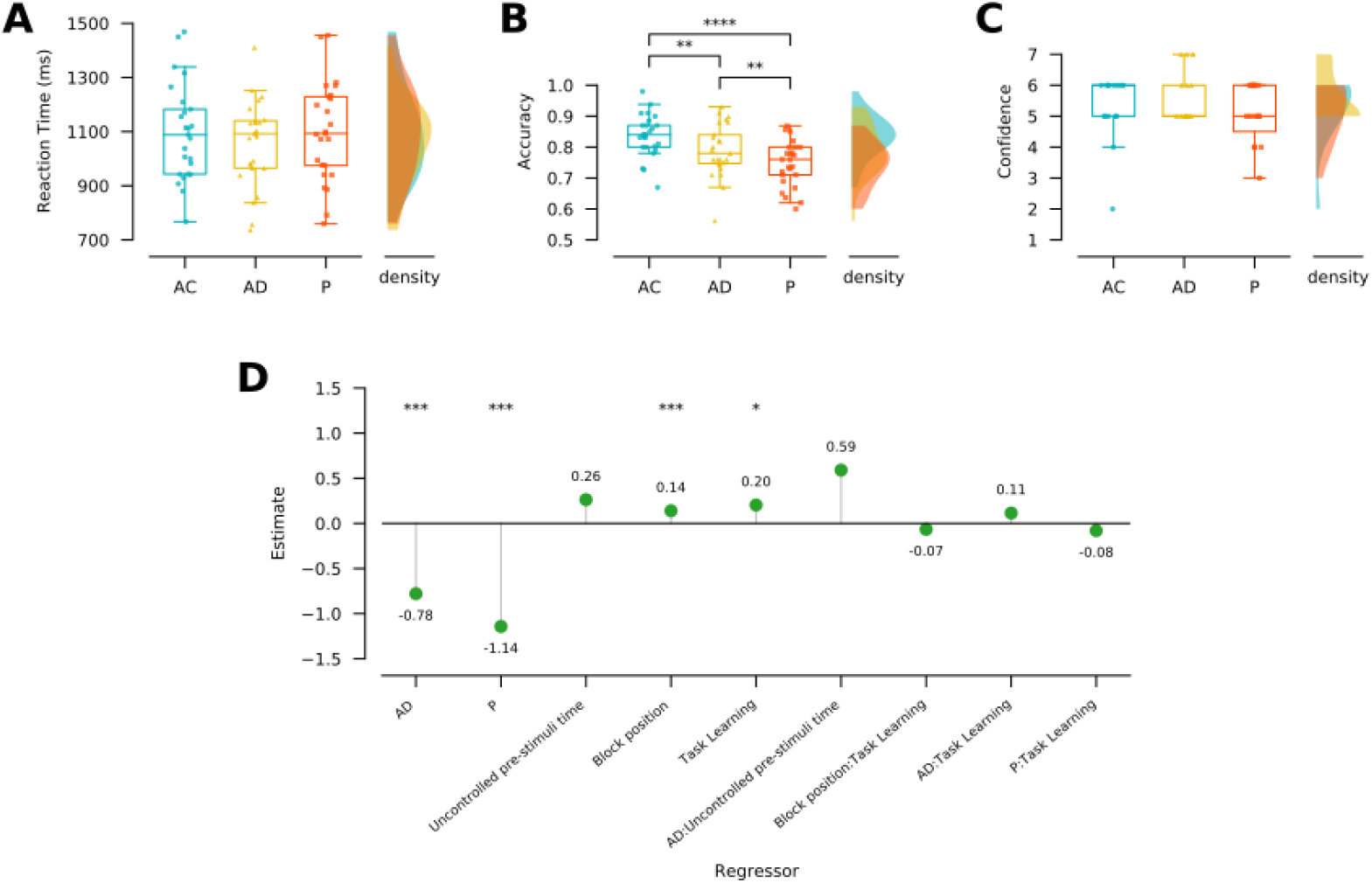
Agents’ self-initiation of the stimuli enhances WM performance. **A. Reaction times** per encoding condition. Left panel, box plot shows the reaction times in milliseconds (ms) per condition (AC: light blue; AD: yellow; P: orange). Right panel, density plot shows the distributions of the reaction times per encoding condition (n=24). Dots in box plots represent the value for each participant. Upper limit of the box = 75^th^ percentile; lower limit of the box = 25^th^ percentile; upper whisker = upper limit value; lower whisker = lower limit value; outlier values are shown outside the whiskers. B. Similar to A, but for accuracy (n=24). C. Similar to A, but for Confidence ratings (n=15). D. Effects of variables (regressor) over the estimated value of making a correct answer. Colon (:) indicates interactions between parameters. (*: p≤ 0.05; **: p ≤ 0.01; ***: p≤ 0.001; ****: p ≤ 0.0001).

To assess if agents’ self-initiation enhances WM performance, we executed a post hoc test, analyzing the accuracy difference between AC (mean = 0.84 ± 0.07) and P conditions (mean = 0.74 ± 0.07). Paired t-tests corrected by the Holm-Sidak method show that self-initiation presents better accuracy than the P condition (t = 8.214; p = 5.9e-08). To further investigate if the timing between motor acts and its sensory consequences is relevant to the agency effect, a post hoc test analyzing the differences between AC and AD conditions (mean = 0.78 ± 0.08) was performed. Paired t-tests corrected by Holm-Sidak method also confirm better accuracy in AC compared to AD condition (t = 3.259; p = 0.0033). Noteworthy, when subjects engage in a passive WM task, their accuracy is poorer when compared to both active conditions (AC compared to P: t = 8.214; p=5.9e-08, AD compared to P: t = 3.565; p=0.0031). This indicates the impact of agency on WM performance, which seems to be partially linked to action/stimuli coupling. A remarkable point is that these accuracy effects do not seem to be explained by differences in the perceived difficulty of the task since there are no significant differences in their reported confidence, rated by the participant at the end of each condition block (*χ*^2^ = 1.472; p = 0.478) (Fig. 2C).

We then explored the impact of other cognitive factors on performance. The other factors tested were task learning (i.e., the index of trial within the block), block position, total pre-stimuli time (i.e., the time between the first fixation onset and the stimuli onset), and uncontrolled prestimuli time (i.e., the time between the first fixation onset and stimuli onset in the passive condition and the time between the button press and the stimuli onset in the AD and AC condition) (for further details, see <u>Behavioral data analysis</u>). To choose which pre-stimuli time has a greater effect on performance, we performed two binomial General Linear Mixed Models (GLMM). Both models just varied in which pre-stimuli time was used as fixed effects, including trial accuracy as the dependent variable and subjects as a random effect. The first model’s fixed effects were encoding condition, task learning, block position, and total pre-stimuli time. The second model’s fixed effects were encoding condition, task learning, block position, and uncontrolled pre-stimuli time. According to the Akaike Information Criterion (AIC), the second model is significantly better than the first model (△ AIC = 7,2). The best model results show a significant effect in three of the independent variables assessed: task learning (***β*** = 0.20±0.09, z = 2.25, p = 0.024), block position (***β*** = 0.139±0.04, z = 3.914, p = 9.1e-05), and the encoding conditions (AD condition: ***β*** = −0.779±0.195, z = −3.992, p = 6.55e-05; P condition: ***β*** = −0.976±0.317, z = −3.077, p = 0.002) (Fig. 2D). Although the second model including the uncontrolled pre-stimuli time was better than the first one, results yield just a non-significant trend for the uncontrolled pre-stimuli time (***β*** = 0.263±0.136, z = 1.923, p = 0.054). This implies that the probability of making a correct response is not modulated by the random time that the stimuli takes to appear in the AD and P conditions. We did not find significant interaction between task learning and block position (***β*** = −0.065±0.03, z = −1.840, p = 0.06), nor interaction between task learning and tasks (Task Learning and AD condition: ***β*** = 0.114±0.07, z = 1.55, p = 0.120; Task Learning and P condition: ***β*** = −0.080±0.719, z = −1.114, p = 0.26). These GLMM results corroborate the previous ANOVA analysis, indicating a significant decrease in the probability of making a correct response when subjects are engaged in the AD or the P conditions compared to AC condition. The probability of making a correct response falls 0.77 log-odds points in the AD condition, and the P condition’s 1.14 log-odds point. Alongside the task learning effect, results show that the probability of making a correct response rises in 0.2 log-odds points with each trial (from 1 to 100) within one block. Results also yield no interaction between task learning and encoding conditions, showing that task learning has the same effect on all three conditions. Likewise, the model reveals that the probability of answering correctly increases in 0.13 log odds with the block position. Like in task learning, the effect of the block position is also independent of the encoding condition. Altogether, GLMM yields other factors such as task learning and block position that influence performance alongside the task conditions effects.

In summary, behavioral analyses show that coupled self-initiation of the stimuli in a WM task increases the probability of performing correctly, even though other timings and learning factors also modulate this probability.

### 2. Agents’ self-initiation of the stimuli accelerate N1 and P2 latencies during WM encoding

We then assessed whether self-initiation of the stimuli impacts visual, attentional, and memory updating processing during WM encoding. To do so, the amplitudes and latencies of related ERP indexes were analyzed: N100-like component, proposed as an index of early visual discrimination related to the visual cortex (Vogel & Luck 2000); P200-like component, which is thought to reflect attentional processing of stimuli in WM (Dunn et al. 1998); and P300-like component, a marker proposed as an index of mental revision of the stimuli (Polich 2007). The ERP’s grand averages for correct trials are shown in Figure 3, specified per encoding conditions.

**Figure 3.**
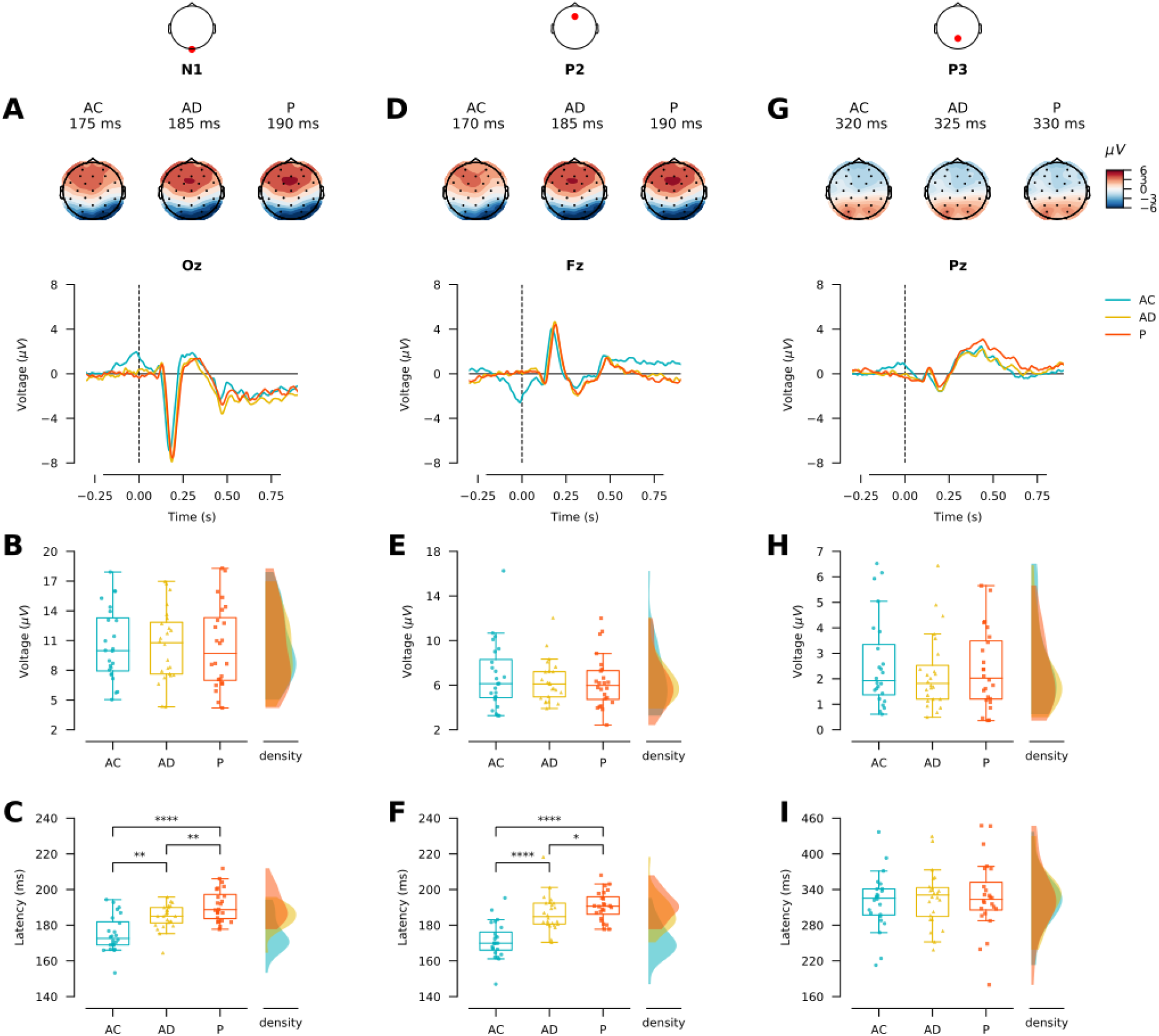
Agents’ self-initiation modulates early visual and attentional ERPs indexes. **A.** Upper panel Topographical plots of the indicated times. Dots represent modeled electrode positions; red: positive voltage; blue: negative voltage; values in μV. Bottom panel. ERP grand average (n = 24 subjects) evoked by stimuli presentation (t = 0 ms) in electrode Oz, per conditions. Only correct trials are included. **B.** Left panel, box plot shows the peak-to-peak voltage (μV) of N1 component, per condition. Right panel, density plot shows the distributions of peak-to-peak voltage of N1 per encoding condition. **C.** Similar to B, but for N1 latency (in ms). **D**. Similar to A, but for electrode Fz. E. Similar to B, but for P2 peak-to-peak amplitude. **F.** Similar to B, but for P2 latency **G.** Similar to A, but for electrode Pz. **H**. Similar to B, but for P3 peak-to-peak amplitude. **I**. Similar to B, but for P3 latency. (*: p ≤ 0.05; **: p ≤ 0.01; ****: p ≤ 0.0001).

The ANOVA results reveal a significant effect of task conditions on the latency of the N1 (F(2,46) = 28.88, p = 7.48e-09; η^2^G = 0.37) and the P2 components (F(2,46) = 46.701, p = 8.418e-12, η^2^G = 0.449). No statistically significant effect of encoding conditions were found on P3 latency (χ^2^ = 0.242, p = 0.886) and on amplitude of all the components analyzed (N1: F(2,46) = 0.037, p = 0.963; P2: *χ*^2^ = 0.089, p = 0.956; P3: *χ*^2^ = 0.328, p=0.848). These results suggest a modulatory effect of task conditions on sensory and attentional processes, but not on memory updating mechanisms.

Two post hoc tests were performed, assessing the difference of N1 and P2 latencies between AC and P conditions. Paired t-tests corrected by the Holm-Sidak method show that subjects have earlier latencies in AC compared to P condition, both in N1 (t = −8.204; p = 8.3e-08) and P2 (t = −9.431; p = 6.8e-09). Passive condition, on the other hand, yields the latest latencies in both N1 (AC/P: t = −8.204; p = 8.3e-08; AD/P: t = −3.009; p = 0.006) and P2 (AC/P: t = −9.431; p = 6.8e-09; AD/P: t = −2.1; p = 0.047) components. These results suggest an effect of self-initiation of the stimuli on sensory and attentional processing.

To further examine if the temporal coupling between motor acts and its sensory consequences is relevant to the agency effect, two post hoc tests we performed, analyzing the N1 and P2 latency differences between AC and AD conditions. Paired t-tests corrected by the Holm-Sidak method also confirm earlier latencies of AC both in N1 (t = −4.224; p = 0.0006) and P2 (t = −6.461; p = p=2.7e-06). As in behavioral analysis, these ERP results suggest that the agency effect is partially linked to the temporal coupling between action and its sensory consequence.

Note that all the three electrodes plotted (Oz, Fz, and Pz) show a component around stimuli presentations, specifically in AC condition. We interpret this component as a neural marker of the motor system activity related to the button press. Consistently, this component is also present in AD condition when ERPs are locked to the button press instead of stimuli presentation (See Figure S1).

In summary, the abovementioned results support that coupled self-initiation modulates the temporal domain of the neural mechanisms underlying the visual discrimination process (N1 component) and the attentional processes engaged during encoding in WM (P2 component), but there is no statistically significant effect on memory updating mechanisms (P3 component). Moreover, no effects on amplitudes were observed. Results also show that delays between movements and their sensory consequences (as reflected in AD condition) do not yield the same effect on encoding processing. Nevertheless, on comparing AD to passive condition, the latencies of N1 and P2 appear earlier.

### 3. Earlier attentional processing is a marker of a coupled active phenomenon

Next, we explored whether the earlier deploying time of visual and/or attentional processing provides reliable markers of self-initiation of stimuli. To do so, we performed a conditional Classification Tree (CART model) that allowed us to test how reliably we can estimate the experimental condition of a trial from the latency of its associated P1 and P2 components. Since CART models are ideal for classifying or evaluating discrete state variables (Strobl et al. 2009), by using this method, it is assumed that initiating the trial in a coupled way would deploy a phenomenon presumably absent during decoupled or passive conditions. CART model results show that the latency of the P2 component better distinguishes between encoding conditions, using a cutoff of 175.293 ms (p < 0.001) (See Fig. 4A). No other ERP variable had a significant effect according to the model. Latencies below 175 ms corresponded mostly to the AC condition (AC median = 169.92 ± 10.14 ms), while later P2 latencies tended to be found in AD (median = 184.81 ± 10.2 ms) and P conditions (median = 190.67 ± 8.09 ms). This result suggests that the attentional index’s onset time is the variable that better distinguishes a coupled self-initiated encoding process from a decoupled or a passive one.

**Figure 4.**
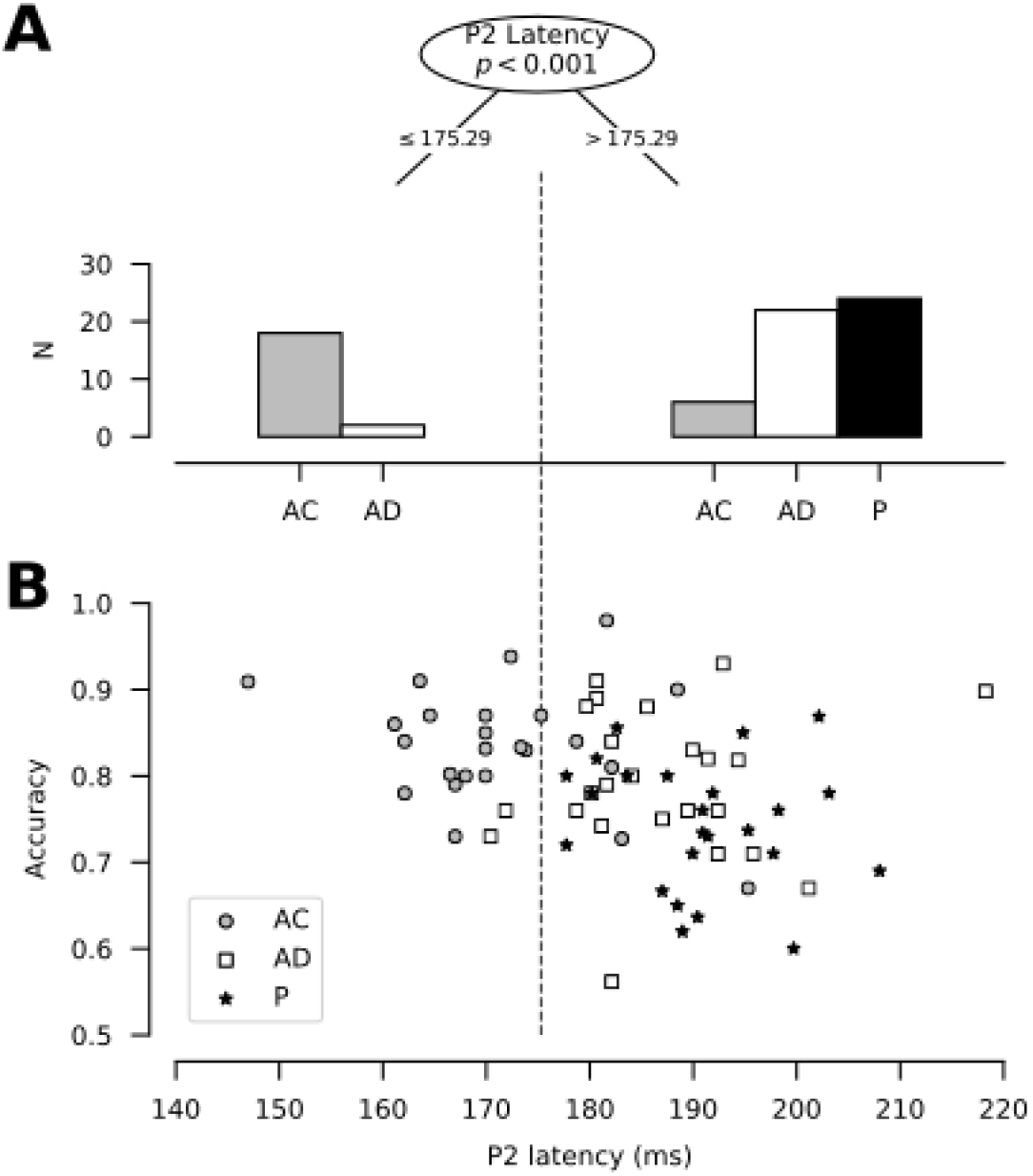
CART model of encoding conditions. **A**. Histograms of the number of participants per encoding condition (y-axis) attributed by the CART node based on P2 latency, with a split at 175.29 ms (top). The left histogram represents latencies lower than the split. Conversely, the right histogram represents latencies larger than the split. The number of cases per condition is equal to 24. **B**. Scatterplot of Accuracy (y-axis) as a function of the latency of the P2 component (x-axis), depicted by the condition (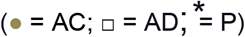 = AC; □ = AD; = P). The discontinuous vertical dashed line represents the split value of the CART model (175.293 ms). Each mark (whether circle, square or star) represents one participant (n = 24 per encoding condition).

Since earlier P2 latency is the variable that better distinguishes the AC condition and that AC is also the condition that shows better accuracy (See Fig. 2B), we analyzed if P2 latency can explain the performance improvement by itself or if it is dependent on the task condition. When contrasted with accuracy, P2 latencies of the AC condition (≤ 175 ms) tend to show higher accuracy values (Fig. 4B). On the other hand, P2 latencies later than 175 ms (mostly compound of AD and P conditions values) tend to show worst accuracy with later latency values (Fig. 4B). To test if this observed relation between P2 and accuracy is dependent on the task condition, we designed a Linear mixed model including accuracy as a dependent variable, P2 latency as a fixed effect, and encoding conditions as random effect variable. Results yield that there is no significant effect of P2 latency on accuracy when including task conditions as random effect variable (***β*** = −0.001 ± 9e-04, t = −1.259, p = 0.106). This result supports that it is not P2 latency by itself that mediates WM improvement.

In summary, the findings show that P2 latency is the variable that better distinguishes a coupled self-initiated encoding process from a decoupled or a passive one, and that it is not P2 latency by itself that mediates WM improvement but the precise nature of the coupled self-initiation condition.

## DISCUSSION

The current study assesses whether the agency modulates WM encoding through temporal modulation of visual and attentional processing. This was investigated by evaluating the influence of active self-initiation of the stimuli on behavior in a Sternberg working memory task, while concurrently measuring ERPs’ encoding components widely used as indexes of visual (N1), attentional (P2) activity, and memory updating mechanisms (P3).

### Agency enhances WM performance and accelerates visual and attentional processing

Our results show that actively initiating the stimuli’s presentation leads to performance enhancement in a WM task. This finding is consistent with previous research stating that WM and motor systems share cognitive resources (Weigelt et al. 2009; Spiegel et al. 2013; Logan & Fischman 2015; Buszard et al. 2016; Gündüz et al. 2017). The present data support that behavioral enhancement is not explained as differences in task difficulty since there is no distinction in the perceived confidence among the three task conditions. Alongside this, there is no statistically significant difference in reaction times (RT), which is classically modulated by task difficulty (Nickerson 1972; Pins & Bonnet 1997; Schneider & Anderson 2011). Our findings rather suggest a role of temporal predictability or motor systems modulation on cognition.

It has been widely proposed that motor actions can predict the timing (i.e., “when”) of the stimulus onset (Saleh et al. 2010; Fujioka et al. 2012; Arnal & Giraud 2012). Motor regions such as the primary sensorimotor cortex (SM1), supplementary motor area (SMA), and cerebellum have been proved to be sensitive to auditory regularities even when auditory stimuli are not under the focus of attention, suggesting a role of these cortices on the temporal prediction of stimuli (Fujioka et al. 2012). While temporal prediction mediated by both overt and covert movements tend to reduce sensation and its neural correlates (Blakemore et al. 1999; Voss et al. 2008; Wolpe et al. 2018; Klaffehn et al. 2019; but for a contradictory effect, see Rohenkohl et al. 2014), no evidence of such reduction in components’ amplitude was found in the current work (See Fig. 3). On the contrary, our results show a temporal modulatory effect on visual processing, which is more in line with previous studies suggesting that motor-mediated temporal prediction facilitates sensorial processing (Ito et al. 2011; Devia, Montefusco-Siegmund, Egaña, Maldonado 2017; Schwarz et al. 2018). On the other hand, when temporal prediction interacts with both bottom-up and topdown attentional orientation mechanisms, time prediction boosts performance and its neural correlates (Kok et al. 2012; Marchant & Driver 2013; Morillon et al. 2016; Kaiser & Schütz-Bosbach 2018). Consistent with these reports, our findings show that time-locked motor-mediated triggering of the stimuli is associated with both attentional acceleration and performance improvement in a WM task.

Furthermore, our results suggest that, although coupled self-initiation of the stimuli is correlated to earlier sensory and attentional processes, the latter mechanism better distinguishes a coupled self-initiated encoding process from a decoupled or a passive one. It is noteworthy that no effects on the amplitudes of N1, P2, or P3 were found, suggesting that self-initiation effects are not based on modulating the number of neural populations required by the task but on influencing the time domain in which encoding is deployed. Altogether, coupled self-initiation of the stimuli enhances WM performance and engages a faster attentional state related to WM encoding, which seems to match the effects of temporal predictability previously reported. Our results support that this attentional state is not engaged during decoupled self-initiation.

If the agency’s effect is solely based on temporal predictability, it should be suppressed under active-unpredictable situations. Our results show that this is not the case: decoupled action-stimuli triggering yields better accuracy and earlier latencies than passive stimuli onset. This result suggests that temporal predictability is not the only mechanism involved in the agency’s WM effect. Embodied cognition theory states that subjects’ bodies, particularly their motor systems, influence cognition (O’Regan & Noë 2001; Varela et al. 2016). In agreement, sensory attenuation effect (i.e., the decreasing in neural sensory response due to self-initiation of a stimulus in a perceptual task) prevails even when compared to an externally generated tone, which is equally predictable in terms of time onset (Klaffehn et al. 2019), suggesting that sensory attenuation effect is not solely due to temporal predictability.

Moreover, decoupled movements modulate perception (Tomassini et al. 2015) and attention (Nittono 2007). There is also evidence suggesting that unrelated button pressing improves long-term memory encoding, whether the action is time-locked to stimulus onset or not (Yebra et al. 2019). Our results corroborate a parallel non-predictive modulation of motor systems over WM. This modulation is possibly related to the recruitment of catecholaminergic pathways, which will be discussed further.

### Agency does not modulate memory updating during WM encoding

It was additionally hypothesized that agency could modulate the P3 component during WM encoding. P3 has been proposed as an index of the mental revision of WM stimuli (Polich 2007), and it seems to be related to posterior parietal cortex activity (Knight et al. 1989; Verleger et al. 1994). Classic WM studies have shown that the amplitude of P3 during encoding is an index of successful encoding, such that greater amplitude of P3 during encoding correlates with later successful retrieval (Karis et al. 1984; Fabiani et al. 1986). Our lack of effect on the P3 component implies that memory updating of stimuli in WM seems to be equal in both active and passive stimuli triggering. It should moreover be noted that the effect reported by Karis et al. (1984) and Fabiani et al. (1986) is dependent on task and rehearsal strategy used by the subjects, such that it is related to salient stimuli and non-elaborative memory rehearsal strategies (i.e., when subjects do not relate stimuli to retain them). These studies report that, even though elaborative rehearsal is associated with better accuracy, the relation between P3 amplitude and accuracy is not further evident when subjects use this strategy.

In our work, subjects did not receive any special instruction about the rehearsal strategy, so it is possible that some of the participants could have used elaborative rehearsal strategies. As the strategy report was not requested, it is not feasible to corroborate if this can explain the lack of effect. On the other hand, our work did not manipulate the stimuli’s saliency, being all consonants of the same size, contrast, and luminance. Future studies could explore the interaction between agency and bottom-up attentional mechanisms.

### The significance of the encoding stage of WM

Notably, the proposal held in this work emphasizes that encoding processing is an equally relevant component of WM, since encoding implies WM mechanisms’ initiation, using perception to memorize, in contrast to the user perception for purely discriminatory purposes. In this sense, as WM mechanisms engage during encoding, self-initiation modulates attentional mechanisms deployed during WM encoding, impacting WM’s overall performance. The WMs’ relationship with the encoding process as part of WM has been extensively explored in the auditory and verbal domain, such that the association between auditory perception—and its deficit—and verbal WM is well documented (e.g., Norrelgen et al. 2002; Wayne et al. 2016; Zhang et al. 2016; Guijo et al. 2018; Magimairaj & Nagaraj 2018). This relationship appears to be less investigated in the visual modality (Cattaneo et al. 2011; Qian et al. 2017; Kosilo et al. 2017). It might be attributable due to the close connection between oral language and WM. One of the central and classic focuses of WM studies is its correlation to oral language, oral language development, and oral language disorders (e.g., Baddeley et al. 1998; Caplan & Waters 1999; Montgomery 2003; Jonsdottir et al. 2005; Coelho et al. 2013; Vugs et al. 2013; Vugs et al. 2014; Fortunato-Tavares et al. 2015; Cogan et al. 2017). In this sense, the relationship between auditory deficits, cognitive effort, and language development appears to be more natural. On the contrary, and to the extent of our review, we did not find further research on visual signal degradation—either by experimental methods, such as luminance or contrast manipulations or by subjects’ impairments, as low vision—on WM, which is relevant to assess due to the critical role of encoding within WM, as our findings have remarked it.

### Neural Model

Based on the results of this study, we argue that the neural mechanisms underlying the time-coupled self-initiation effect on WM might be based on motor cortex modulation on perceptual and attentional neural systems. Motor cortices, such as M2 in rats or the premotor cortex in primates, are anatomically connected to the visual cortex (Attinger et al. 2017; Leinweber et al. 2017). Some theories propose that motor-sensory cortex projections may act as efference copies used by an internal forward model to predict the sensory consequence of the motor act (Miall & Wolpert 1996; Wolpert & Kawato 1998). A proposed role for these projections is the sense of agency, allowing the nervous system to discriminate when a sensory consequence is externally or self-generated (Poletti et al. 2017). Another role for motor-sensory cortex projections is to generate time predictions about the onset of sensory consequences, acting as a marker about when a sensory change will happen (Saleh et al. 2010; Arnal & Giraud 2012). This would possibly occur through the modulation of local field potentials in the sensory cortex, as shown in the visual cortex (Ito et al. 2011). In our data, N100 lower latencies in time-coupled self-initiated conditions reflex early visual processing facilitation (Schroeder et al. 2010). This temporal modulation could be due to motor cortices activating motor-sensory cortex projections. This hypothesis should be tested in further studies, including methods for localizing the source of the electroencephalographic activity (which requires a greater number of recording channels) or magnetoencephalography.

Likewise, P2 earlier latencies in coupled self-initiated conditions can reflect the attentional processing’s facilitation during WM encoding originated from the same motor cortices signaling. Our results suggest that this mechanism could operate in non-sensory cortex related to later attentional processing important to WM such as the dorsolateral prefrontal cortex (DLPFC) (Blumenfeld & Ranganath 2006; D’Esposito & Postle 2015) and posterior parietal cortex (PPC) (Todd & Marois 2005; Curtis 2006; Berryhill & Olson 2008). Motor cortices project to both DLPFC (Hasan et al. 2013) and PPC (Reep et al. 1994; Wilber et al. 2015), probably via the superior longitudinal fascicle connecting the supplementary motor cortex with the abovementioned cortices (Bozkurt et al. 2017). Therefore, movement-related activation of SMA might be signaling the income of a relevant stimulus to be maintained to DLPFC and PPC cortices as well. Consequently, this signal may facilitate the activation of the frontoparietal network underlying attention, such that this network might have a faster activation when it receives a motor signaling. Since cortico-cortical activations through direct anatomic projections occur within a few milliseconds (Leinweber et al. 2017), this effect should be visible shortly after the activation of the motor cortex. Hence, the temporality between the movement and the relevant stimuli presentation seems to be a relevant feature of the modulatory mechanism of the active self-initiation of the stimuli, as supported by previous studies (Blakemore et al. 1999; Morillon et al. 2014; Concha-Miranda et al. 2019) and our results. Notably, this appears not to affect the memory updating of WM stimuli, as our P3 results suggest, even though this component has possibly originated in the PPC cortex (Knight et al. 1989; Verleger et al. 1994).

Remarkably, the motor system can modulate cognition not only using direct corticocortical connections but also through modulatory effects of motor activation on dopamine, acetylcholine, and noradrenergic circuits. The dopamine system’s importance in WM functioning has been extensively documented (Brozoski et al. 1979; Sawaguchi & Goldman-Rakic 1991; Müller et al. 1998; Sawaguchi 2001). For instance, Müller et al. (1998) report a performance enhancement in a WM task when primates were administered with a D1/D2 dopamine agonist, possibly modulating DLPFC through the mesocortical pathway. On the other hand, dopamine depletion in monkeys’ DLPFC induces deficits in visual WM tasks (Brozoski et al. 1979). Strikingly, the same authors demonstrate the reversion of the deficits when monkeys were administered with dopaminergic agonists such as L-dopa and apomorphine. Alongside this, it is known that the ventral tegmental area (VTA, which is related to the mesocortical circuit) is a signal target of motor cortices (Beier et al. 2015). Therefore, a voluntary movement could modulate dopaminergic activity, modulating DLPFC activity during active self-initiated WM encoding. Likewise, cholinergic neurons located in the basal prosencephalon show fast activity related to the current movement (Pinto et al. 2013; Eggermann et al. 2014; Nelson & Mooney 2016). In turn, basal prosencephalon cholinergic neurons projects to DLPFC (Gielow & Zaborszky 2017) and medial prefrontal cortex (Bloem et al. 2014) as well as to sensory cortices (Pinto et al. 2013; Eggermann et al. 2014; Nelson & Mooney 2016). According to this view, a motor action could modulate movement-responsive basal prosencephalon cholinergic neurons, which in turn may modulate prefrontal and sensory cortices during active WM encoding. Finally, evidence shows locus coeruleus adrenergic neurons respond to voluntary movements (Kalwani et al. 2014; Bouret & Richmond 2015). The adrenergic system has been correlated to enhancing long-term memory encoding (Cahill et al. 1994; Yebra et al. 2019). Since long-term memory is postulated as an important component of WM (Cowan 2005), it is possible that it could also participate in active self-initiated WM encoding. It is also known that basal ganglia seem to participate in sensorimotor, associative, and limbic information (Leisman & Melillo 2013; Jahanshahi et al. 2015). As part of the associative circuit functioning, basal ganglia seem to have a supporting role in WM (Constantinidis & Procyk 2004; Marvel et al. 2019). Previous studies report projections between the associative circuit of basal ganglia and the prefrontal cortex (Dum & Strick 2009), specifically between DLPFC and internal globus pallidus-substantia nigra pars reticulata. Although basal ganglia are organized in three different topographically separated circuits, these circuits also overlap, allowing for the integration of associative, sensorimotor, and limbic signaling (Draganski et al. 2008). Thus, the activation of sensorimotor basal ganglia circuits due to a voluntary movement could influence associative basal ganglia circuits and possibly modulate WM.

Critically, the networks mentioned above require several synaptic relays to take place and thus could unravel more slowly than the modulatory effects of direct motor-sensory/associative cortices projections. Accordingly, the catecholamine system’s recruitment and basal ganglia circuits by motor systems during active self-initiated encoding could explain the discrepancy between decoupled self-initiated encoding and passive encoding in both behavior and electrophysiology. On the other hand, fast direct cortico-cortical projections can explain the distinction between time-coupled and decoupled self-initiated encoding. While stimuli appearing coupled to the movement would be modulated by both fast (direct motor projections) and slow (activity related to catecholamines and basal ganglia circuits) modulatory networks, stimuli presented hundreds of milliseconds after the movement has been executed would be influenced by the slow response modulatory networks only. This would explain that both coupled and decoupled self-presentation lead to improved WM performance but that the effect is stronger in the coupled condition.

### Coda

To conclude, the current findings show that agency, present during active self-initiation of the stimuli, modulates WM encoding processing through the influence of early sensory and attentional processes. Performance enhancement in coupled self-initiation is related to an earlier attentional state, which seems to be absent in passive and active decoupled self-initiated states. Nevertheless, processing facilitation of early sensory and attentional processes is also present in decoupled self-initiation compared to the stimuli’s passive triggering. This suggests that active self-initiation, regardless of whether it is time-locked or not, engages a motor-mediated modulation on cognition in addition to the temporal predictive mechanism (the latter being absent in the decoupled self-initiated condition). Finally, our study also remarks that sensory and attentional processing during encoding is a crucial component of WM, emphasizing that WM does not merely maintain the mechanism of absent stimuli.

## Acknowledgments

Special thanks are extended to Dr. Carolina Delgado, Dr. Marcela Peña, and Dr. Rómulo Fuentes for their valuable and constructive suggestions during the planning and development of this research work. We also extend our thanks to Mariana Mella for her assistance with the data acquisition, and to all the staff of Laboratorio de Neurosistemas.

## Notes

Conflict of Interest: None declared.

