## Supplementary Figure for "Agency improves working memory and accelerates visual and attentional processing"

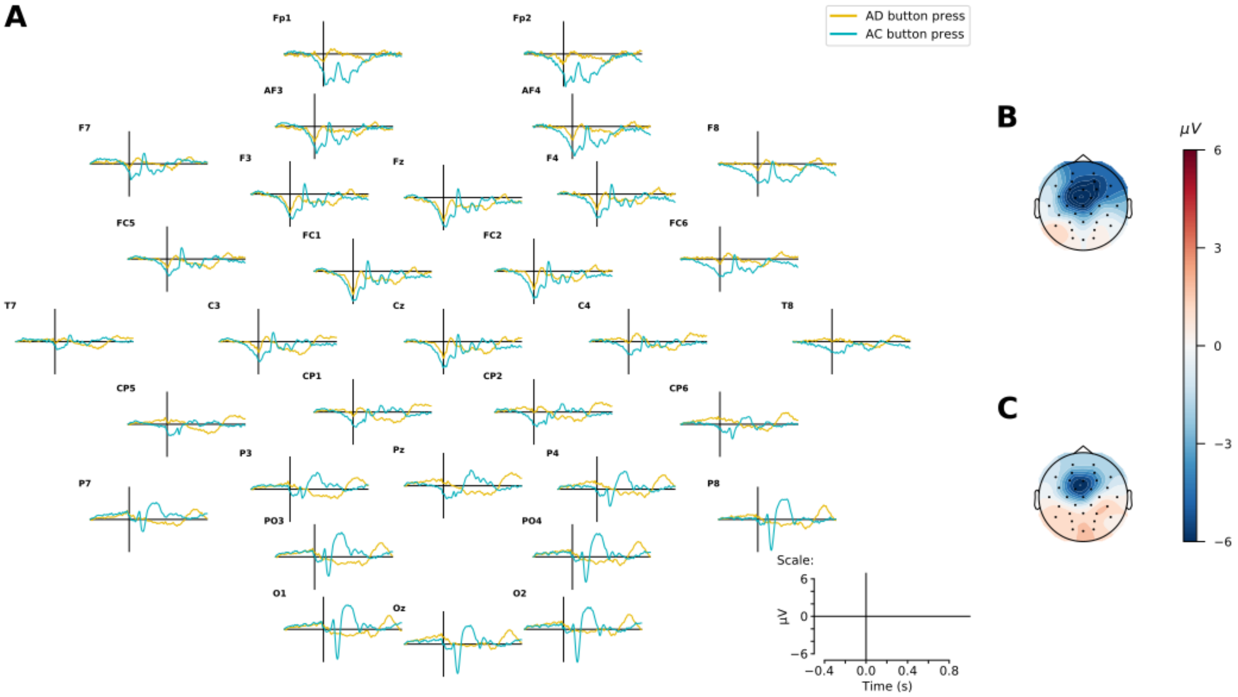


**Figure S1. Grand averages evoked by button press (t = 0 ms). A.** Grand averages ERPs evoked by button press are depicted for all 32 channels. AC is depicted in light blue line, and AD is depicted in yellow line (n = 24 subjects). **B**. Topographical plots at t = 0 ms (button press) for AC condition. red: positive voltage; blue: negative voltage; values in μV. **C.** Same as B, but for AD condition.
